# In vivo glucoregulation and tissue-specific glucose uptake in female Akt substrate 160 kDa knockout rats

**DOI:** 10.1101/777284

**Authors:** Xiaohua Zheng, Edward B. Arias, Nathan R. Qi, Thomas L. Saunders, Gregory D. Cartee

## Abstract

The Rab GTPase activating protein known as Akt substrate of 160 kDa (AS160 or TBC1D4) regulates insulin-stimulated glucose uptake in skeletal muscle, the heart, and white adipose tissue (WAT). A novel rat AS160-knockout (AS160-KO) was created with CRISPR/Cas9 technology. Because female AS160-KO versus wild type (WT) rats had not been previously evaluated, the primary objective of this study was to compare female AS160-KO rats with WT controls for multiple, important metabolism-related endpoints. Body mass and composition, physical activity, and energy expenditure were not different between genotypes. AS160-KO versus WT rats were glucose intolerant based on an oral glucose tolerance test (P<0.001) and insulin resistant based on a hyperinsulinemic-euglycemic clamp (HEC; P<0.001). Tissue glucose uptake during the HEC of female AS160-KO versus WT rats was: 1) significantly lower in epitrochlearis (P<0.05) and extensor digitorum longus (EDL; P<0.01) muscles of AS160-KO compared to WT rats; 2) not different in soleus, gastrocnemius or WAT; and 3) ∼3-fold greater in the heart (P<0.05). GLUT4 protein content was reduced in AS160-KO versus WT rats in the epitrochlearis (P<0.05), EDL (P<0.05), gastrocnemius (P<0.05), soleus (P<0.05), WAT (P<0.05), and the heart (P<0.005). Insulin-stimulated glucose uptake by isolated epitrochlearis and soleus muscles was lower (P<0.001) in AS160-KO versus WT rats. Akt phosphorylation of insulin-stimulated tissues was not different between the genotypes. A secondary objective was to probe processes that might account for the genotype-related increase in myocardial glucose uptake, including glucose transporter protein abundance (GLUT1, GLUT4, GLUT8, SGLT1), hexokinase II protein abundance, and stimulation of the AMP-activated protein kinase (AMPK) pathway. None of these parameters differed between genotypes. Metabolic phenotyping in the current study revealed AS160 deficiency produced a profound glucoregulatory phenotype in female AS160-KO rats that was strikingly similar to the results previously reported in male AS160-KO rats.

## Introduction

The Rab GTPase activating protein known as Akt substrate of 160 kDa (also known as AS160 or TBC1D4) is highly expressed by multiple tissues, including skeletal muscle, the heart, and white adipose tissue (WAT) [1–5]. These tissues are important sites for insulin-mediated glucose disposal, and phosphosite-specific phosphorylation of AS160 by Akt is crucial for insulin-stimulated GLUT4 glucose transporter exocytosis and enhanced glucose transport. Accordingly, understanding the relationship between AS160 and glucose uptake in these tissues has implications for whole body glucoregulation and insulin sensitivity.

AS160 deficiency in humans [6], mice [3, 4] and rats [5] results in whole body insulin resistance, but there is limited knowledge about the effects of AS160 deficiency in females, regardless of species. Published research in humans has not addressed the possibility that AS160 deficiency might not have identical consequences on males versus females [6]. Several studies using AS160-KO mice reported data only [4, 7] or mostly [8] in males. Other studies reported data for both male and female AS160-KO mice for some, but not all outcomes [3, 9]. Hyperinsulinemic-euglycemic clamps (HEC) have been performed only in male AS160-KO mice [4], and in vivo tissue-specific glucose uptake has been reported in male, but not female, mice [3, 9]. Currently available research in mice indicates that the metabolic phenotypes of male and female AS160-KO mice are very similar, but not identical. For example, in vivo insulin resistance (based on an insulin tolerance test) was evident in both male and female AS160-KO mice [3]. However, glucose tolerance was normal for male, but not female AS160-KO mice [3]. The only previous study of AS160-KO rats focused exclusively on male animals [5].

Others have independently generated AS160-KO mice [3, 7–9]. However, there is unique value in creating and characterizing a preclinical model in a second species. Because rats are much larger than mice, studying rats offers significant benefits with regard to performing certain surgical procedures and providing substantially greater amounts of tissue to be analyzed [10]. A method was recently described using rat skeletal muscle that enables the measurement of glucose uptake and fiber type based on myosin heavy chain isoform expression in a single muscle fiber [11, 12], but it is unlikely these analyses would be feasible using single fibers from mouse skeletal muscle. Rats have also proven to be extremely valuable for detecting a potential role for AS160 in the processes responsible for improved insulin sensitivity in skeletal muscle [13–16], so information about both male and female AS160-KO rats will be useful for future mechanistic studies.

Male AS160-KO compared to wild type (WT) rats were characterized by lower glucose uptake in insulin-stimulated skeletal muscles [5]. Insulin resistance was accompanied by unaltered proximal insulin signaling as reflected by Akt phosphorylation in skeletal muscle. Consistent with results observed for skeletal muscle of male AS160-KO mice [3, 4, 7], GLUT4 protein abundance was lower in several skeletal muscles from male AS160-KO versus WT rats. GLUT4 abundance was lower in the soleus muscle from female AS160-KO compared to WT mice [3]. In addition to insulin resistance, skeletal muscle from male AS160-KO compared to WT rats [5] and mice [7] had reduced glucose uptake in response to AICAR, which stimulates AMP-activated protein kinase (AMPK), concomitant with unaltered AMPK phosphorylation.

An unexpected discovery in male AS160-KO rats was that myocardial glucose uptake during an HEC was substantially increased compared to WT controls even though GLUT4 protein abundance was lower in the AS160-KO rats [5]. To assess potential mechanisms that might account for the elevated myocardial glucose uptake, we evaluated heart abundance of GLUT1 and SGLT1 glucose transporters, hexokinase II and the SERCA2 calcium pump in male AS160-KO rats. However no genotype differences were evident for any of these proteins in male rats.

Male and female AS160-KO rats cannot be assumed to have identical metabolic phenotypes. Therefore, the primary aim of the present study was to provide the first comparison of female AS160-KO rats and WT controls for a number of important measurements, including: body composition, energy expenditure, food intake, physical activity, glucose tolerance, whole body insulin sensitivity during an HEC, in vivo tissue-specific glucose uptake (skeletal muscle, WAT and heart), hepatic glucose production under basal and insulin-stimulated conditions, and glucose uptake by isolated skeletal muscles in response to insulin and AICAR. As in the earlier study with male AS160-KO rats, protein abundance of GLUT1, GLUT4, hexokinase II, TBC1D1 (an AS160 paralog), and Akt2 phosphorylation were also determined.

The second major aim of the present study was to advance understanding related to our discovery that myocardial glucose uptake was markedly increased in AS160-KO compared to WT rats. In this context, in addition to the analyses performed in the earlier study, we also evaluated other possible mechanisms, including the role of the AMPK pathway (by evaluating phosphorylation of key regulatory sites on AMPK and its substrates acetyl Co-A carboxylase and TBC1D1), the abundance of GLUT8 glucose transporter protein, and the abundance of important metabolic proteins expressed by the heart (lactate dehydrogenase, and multiple components of the electron transport chain and oxidative phosphorylation).

## Materials and methods

### Materials

The apparatus and reagents used for SDS-PAGE and immunoblotting were from Bio-Rad Laboratories (Hercules, CA). T-PER tissue protein extraction reagent (#78510), bicinchoninic acid protein assay (#23225) and MemCode Reversible Protein Stain Kit (#24585) were purchased from Thermo Fisher Scientific (Waltham, MA). Human recombinant insulin was from Eli Lilly (Indianapolis, IN). [^3^H]-2-Deoxy-D-glucose ([^3^H]-2-DG), [1-^14^C]-2-deoxyglucose ([^14^C]-2-DG) and [^14^C]-mannitol were from PerkinElmer (Boston, MA). Mouse/rat insulin ELISA kit (#EZRMI-13K) was from Millipore Sigma (Burlington, MA). Non-esterified fatty acid (NEFA) colorimetric kit (HR Series NEFA-HR) was from Wako Diagnostics (Mountain View, CA). Anti-AS160 (#ABS54), pTBC1D1^Ser237^ (#07-2268) and anti-GLUT4 (#CBL243) were from EMD Millipore (Burlington, MA). Anti-phospho AS160 Thr^642^ (pAS160^Thr642^, #8881), anti-Akt (#4691), anti-phospho Akt Thr^308^ (pAkt^Thr308^; #13038), anti-phospho Akt Ser^473^ (pAkt^Ser473^; #4060), anti-Hexokinase II (#2867), anti-AMP-activated protein kinase (AMPK, #2532), anti-phospho AMPK Thr^172^ (pAMPK^Thr172^, #2535), anti-TBC1D1 (#4629), anti-phospho Akt2^Ser474^ (pAkt2 Ser474, #8599), anti-SGLT1 (#5042), ACC (#3676), pACC^Ser79^ (#3661), anti-LDH (#3558), and anti-rabbit IgG horseradish peroxidase conjugate (#7074) were from Cell Signaling Technology (Danvers, MA). Anti-GLUT1 (#sc-1603) and anti-CD36 (#sc-13572) were from Santa Cruz Biotechnology (Santa Cruz, CA). Anti-SERCA2 ATPase (#S1314) was from Sigma-Aldrich. GLUT8 antibody was generously provided by Dr. Jeremie Ferey from the Washington University School of Medicine in St. Louis. OXPHOS Rodent WB Antibody Cocktail (#ab110413) from Abcam (Cambridge, MA) contains antibodies recognizing the following five proteins from the electron transport chain and oxidative phosphorylation: Complex 1 component, NADH dehydrogenase (ubiquinone) 1β subcomplex subunit 8 (NDUFB8); Complex II component, succinate dehydrogenase complex subunit 8 (SDHB); Complex III component, ubiquinol-cytochrome-c reductase complex core protein 2 (UQCRC2); Complex IV component, Cytochrome c oxidase subunit I (MTCO1); and Complex V component, mitochondrial membrane ATP synthase (ATP5A). All other reagents were either from Fisher Scientific (Hanover Park, IL) or Sigma-Aldrich (St. Louis, MO).

### Animal treatment

The University of Michigan Committee on Care and Use of Animals approved the animal care and breeding procedures, which were performed according to the Guide for the Care and Use of Laboratory Animals from the National Research Council. Animals were housed with controlled lighting (12 h light/dark cycle: lights on at 0500/lights off at 1700) and temperature (22°C) and ad libitum access to rodent chow (Laboratory Diet no. 5L0D; Lab Diet, St. Louis, MO) until an overnight fast at 1700 on the night prior to the terminal experiments.

### AS160 mutant rats created with CRISPR/Cas9

A genetically modified rat line with an *AS160 (TBC1D4)* gene knockout was created on the Wistar outbred genetic background using CRISPR/Cas9 technology [17, 18] as previously described [5]. The insertion of a premature termination codon in exon 1 is predicted to result in loss of protein expression due to nonsense medicated decay of mRNA [19]. A single guide RNA (sgRNA) and protospacer adjacent motif was designed targeting coding strand: 5’ GCGACAAGCGCTTCCGGCTA TGG 3’ with a predicted cut site 111 bp downstream of the initiation codon. The sgRNA target was cloned into plasmid pX330 (Addgene.org plasmid #42230, a generous gift from Feng Zhang) as previously described [20].

*AS160* exon 1 targeting was assessed using electroporation of Sprague Dawley (SD) rat embryonic fibroblasts followed by PCR amplification of the target sequence and analysis for indel formation by T7 endonuclease 1 digestion as described [21]. The circular pX330 plasmid containing the active *AS160* exon 1 target (5 ng/µl final concentration) [22] was used for pronuclear microinjection of rat zygotes. Fertilized eggs for microinjection were produced by mating superovulated Wistar female rats (Envigo Hsd:WI, Strain Code 001) with Wistar males. Pronuclear microinjection was performed as previously described [23]. A total of 297 zygotes were microinjected, 266 surviving zygotes were transferred to pseudopregnant SD female rats (Charles River Laboratory, SAS SD, Strain Code 400), and 58 pups were born. Genotyping of genomic DNA extracted from tail tip biopsies using PCR identified 24 founder rats with indels in *AS160* exon 1.

### Rat breeding and genotyping

Animals were genotyped using PCR with DNA isolated from tail tip samples with primers flanking the sgRNA binding site: 5′-GGCTGGTGGCACCGAGTCAGG-3′ (forward), 5′-CCGACGGATCTCGGCCATGAG-3′ (reverse) followed by sequencing using a nested primer 5′-GCGCGGTGCCCTCGCTAGGC-3′. A founder line carrying a 20-bp substitution deletion (TGGCGACAAGCGTTCCGGC) was selected for colony expansion and backcrossed with wild type Wistar outbred rats (Charles River Laboratory; Wilmington, VA) to establish an *AS160****^+/-^*** colony. Genotyping was performed by Transnetyx (Cordova, TN) using a forward primer 5′-CCTAGCGCAGCCAGGTG-3′ and reverse primer 5′-TCCTGCGATCCAAGCAAGAC-3′ together with reporters 5′-CCGGAAGCGCTTGTC-3′ and 5′-CCACGTACCATAGGCTTG-3′ to detect WT and mutant, respectively.

Transgenic rats were backcrossed using a WT Wistar [Crl:WI] background (Charles River Laboratories) four times. The transgenic breeding colony was managed by the University of Michigan Unit for Laboratory Animal Medicine (ULAM) husbandry services. Tail tip samples from rats aged 2 weeks were used for genotyping. After weaning, homozygous mutant (AS160-KO) and control WT sibling rats were housed in a ULAM holding facility with a 12 h light/dark cycle with ad libitum access to water and food (5LOD chow).

### Body mass, body composition and tissue masses

Body composition was determined using an NMR analyzer (Minispec LF9011, Bruker Optics), and body mass was assessed in rats at 7-8 weeks-old (n = 6 for each genotype). Another cohort of rats was anesthetized (intraperitoneal injection of ketamine/xylazine cocktail, 50 mg/kg ketamine and 5 mg/kg xylazine) at 11 weeks-old, and body mass was measured in these rats. Skeletal muscles (extensor digitorum longus, EDL, epitrochlearis, gastrocnemius, and soleus), WAT, and the heart from these animals were sampled and weighed.

### Indirect calorimetry, physical activity and food intake

Indirect calorimetry and physical activity were analyzed in rats (n = 6 for each genotype) aged 9 weeks with the Comprehensive Lab Animal Monitoring System (CLAMS, Columbus Instruments, Columbus, OH). Animals were housed in the CLAMS unit for 72 hours with unlimited access to food and water. Food consumption was also measured in the animals at 10-11 weeks-old.

### Oral glucose tolerance test (OGTT)

After a 15-16 hour overnight fast, WT (n = 6) and AS160-KO (n=6) animals (aged 11 weeks) were provided 50% glucose via oral gavage (2.0g/kg). Blood was sampled before and following the gavage (0, 15, 30, 60, and 120 minutes) by the tail vein. Blood glucose was determined with a glucometer (Accu-Chek, Roche), and plasma insulin concentration was assessed by ELISA. The rats were briefly restrained (<1 minute) for each blood collection. Area under the curve (AUC) for glucose and insulin was determined with the trapezoidal rule [24].

### Hyperinsulinemic-euglycemic clamp (HEC)

WT (n = 6) and AS160-KO (n = 6) rats had catheters surgically placed in their jugular vein and carotid artery one week before a HEC experiment conducted when rats were 10 weeks-old as described earlier [5, 25]. At approximately 1700 h on the day before the HEC was performed, food was removed from rat cages (approximately 16 hours prior to the start of the clamp). The protocol included a 90 minute tracer equilibration period (t = −90 to 0 minutes) beginning at ∼0930 h, followed by a 120 minute experimental period (t = 0 to 120 minutes) beginning at ∼1100 h. At t = −90, a bolus infusion of 12 µCi of [3-^3^H]-glucose (HPLC purified; PerkinElmer) was provided, followed by a 0.125 µCi/minute infusion for 90 minutes. At t = −10 minutes, blood was sampled (∼100 µl) to determine basal levels of insulin and glucose and glucose turnover. The insulin infusion initiated at t = 0 began with a primed-continuous infusion (200 mU/kg bolus, followed by 20 mU/kg/minute or 120 pmol/kg/minute) of human insulin (Novo Nordisk). The infusion of [3-^3^H]-glucose was elevated to 0.20 µCi/minute for the remainder of the experiment to minimize alterations in specific activity. Euglycemia (120-130 mg/dL) was maintained by measuring blood glucose every 10 minutes with an Accu-Chek glucometer (Roche, Germany) beginning at t = 0 and infusing 50% glucose at variable rates as needed. Blood was sampled (100 µl) during a steady-state of glucose infusion at t = 80, 85, 90, 100, 110 and 120 minutes for measurement of glucose specific activity. Insulin levels were measured in samples taken at t = −10 and 120 minutes. To estimate tissue glucose uptake, a bolus intravenous injection of [^14^C]-2-DG was given at t=78 minutes while continuously maintaining the HEC steady state. Blood was collected at 2, 7, 12, 22, 32 and 42 minutes after the injection for determination of plasma [^14^C]-2-DG radioactivity. At the end of the experiment, rats were anesthetized with an intravenous infusion of sodium pentobarbital (SP). Tissues [epitrochlearis, gastrocnemius, soleus, EDL, WAT (peri-uterus/fallopian tubes depot), and heart] were rapidly dissected and freeze-clamped with aluminum tongs cooled in liquid N_2_ and stored at −80°C until subsequent processing. A part of the tissues was used to measure 2-DG uptake as previously described [26] and another part of the tissue was used for immunoblotting as described below.

### Plasma insulin and non-esterified fatty acids (NEFA)

Plasma was sampled at −10 and 120 minutes during the HEC and used to measure plasma insulin via ELISA. Plasma samples from −10, 80, 90 and 120 minutes were used to measure NEFA levels with a colorimetric assay.

### Isolated skeletal muscle procedures

At 1600 to 1700 on the night preceding the isolated skeletal muscle procedures, food was removed from the cages of the rats (aged 9 weeks). Rats (n = 8-9) used for insulin treatment experiments were anesthetized with intraperitoneal injections of SP (50 mg/kg), and rats used for the 5-aminoimidazole-4-carboxamide-1-β-D-ribofuranoside (AICAR) experiment were anesthetized with a ketamine (50 mg/kg)/xylazine (5 mg/kg) cocktail (K/X). In preliminary experiments, we found no significant difference for glucose uptake by isolated muscles isolated from animals anesthetized with SP compared to K/X (not shown). For insulin experiments, in animals that were deeply anesthetized, both of their epitrochlearis and soleus muscles were excised. Soleus muscles were longitudinally split into two strips of similar size. Epitrochlearis muscles and soleus muscle strips were placed in vials including the appropriate solution with continuous shaking (50 revolutions per minute) and gassing (95% O_2_ - 5% CO_2_) in a heated (35°C) water bath. Muscles were incubated in vials including 2 ml Krebs Henseleit Buffer (KHB) along with bovine serum albumin (BSA; 0.1%), 2 mM sodium pyruvate, 6 mM mannitol ±insulin (500 µU/ml) for 30 minutes. Following the initial incubation step, muscles were transferred to another vial containing KHB-BSA with the same insulin dose as the first step with 1 mM 2-DG (specific activity of 2.25 mCi/mmol [^3^H]-2-DG), and 9 mM mannitol (specific activity of 0.022 mCi/mmol [^14^C]-mannitol) for 20 minutes. After this incubation step, muscles were quickly blotted on filter paper moistened with ice-cold KHB, trimmed, frozen with aluminum tongs cooled in liquid N_2_, and stored at −80°C until subsequent processing and analysis. The same incubation procedure was employed for experiments using AICAR (n = 6-9), except only the epitrochlearis was studied because previous research indicated that AICAR increased glucose uptake by rat epitrochlearis, but not rat soleus muscle [27, 28].

### Tissue lysate preparation

A portion of the tissues sampled from animals after the clamp was weighed and processed for measurement of [^14^C]-2-DG accumulation as previously described [26]. Another portion from these tissues and the muscles from the ex vivo incubation experiments were weighed, transferred to pre-chilled glass tissue grinding tubes (Kontes, Vineland, NJ), and homogenized in ice-cold lysis buffer (1 ml) with a glass pestle attached to a motorized homogenizer (Caframo, Georgian Bluffs, ON). The lysis buffer included TPER supplemented with 1 mM EDTA, 1 mM EGTA, 2.5 mM sodium pyrophosphate, 1 mM sodium vanadate, 1 mM ß-glycerophosphate, 1 µg/ml leupeptin, and 1 mM phenylmethylsulfonyl fluoride. Homogenates were rotated for 1 h (4°C) before centrifugation (15,000 × g for 15 minutes at 4°C). The supernatants were transferred to microfuge tubes and stored at −80°C until later analyses. Protein concentration was determined by the bicinchoninic acid procedure [29].

### Glucose uptake by isolated muscles

Aliquots (200 μl) of the supernatants from centrifuged lysates that were prepared from incubated muscles were pipetted into a vial with 8 ml of scintillation cocktail (Research Products International, Mount Prospect, IL). A scintillation counter (Perkin Elmer) was used to quantify ^3^H and ^14^C disintegrations per minute. These values were used to calculate [^3^H]-2-DG uptake according to previously described procedures [30, 31].

### Immunoblotting

An equal amount of protein from each sample was combined with Laemmli buffer, boiled (5 minutes) and separated via SDS-PAGE (9% resolving gel) before being transferred to polyvinylidene fluoride (PVDF) membranes. Membranes were blocked (5% BSA or non-fat milk in TBST for 1 hour at room temperature) and then incubated overnight (4°C) with 5% BSA or milk-TBST and the appropriate primary antibody. Membranes underwent 3 × 5 minutes washes in TBST and were then incubated with secondary antibody in 5% BSA or nonfat milk for 1 hour at room temperature. Blots were again washed 3 × 5 minutes in TBST and 2 × 5 minutes TBS. Finally, PVDF membranes were subjected to enhanced chemiluminescence (ECL) to visualize protein bands and immunoreactive proteins were quantified by densitometry (Fluorchem E and AlphaView Imaging Software; ProteinSimple, San Jose, CA). Memcode staining was used for the loading control (sample values were divided by their respective Memcode values), and individual sample values were normalized to the mean of all sample values on each blot.

### Skeletal muscle myosin heavy chain isoform determination

Soleus, EDL, epitrochlearis and gastrocnemius muscles of rats (10 weeks-old; n = 6) were isolated, weighed and processed as previously described for MHC isoform abundance [32]. MHC bands were quantified using densitometry.

### Statistical analysis

Data were statistically analyzed with SigmaPlot 13.0 (San Jose, CA). Comparisons between the WT and AS16O-KO rats were performed with an unpaired two-tailed *t*-test. Comparisons between contralateral muscles that were incubated ex vivo (either ±insulin or ±AICAR) were analyzed using a paired two-tailed *t*-test. Two-way ANOVA was performed to compare more than two groups (ex vivo muscles incubated ±insulin or ±AICAR), and Holm-Sidak post hoc tests were used to identify the source of significant variance. Data were expressed as means ± SEM. P-values ≤0.05 were considered statistically significant.

## Results

### Genotype confirmation

Genotype was determined using qPCR with DNA from tail tip samples. Lack of AS160 in tissues from AS160-KO rats was also substantiated via immunoblotting for AS160 protein after the animals were euthanized (Figure 1).

**Figure 1.**
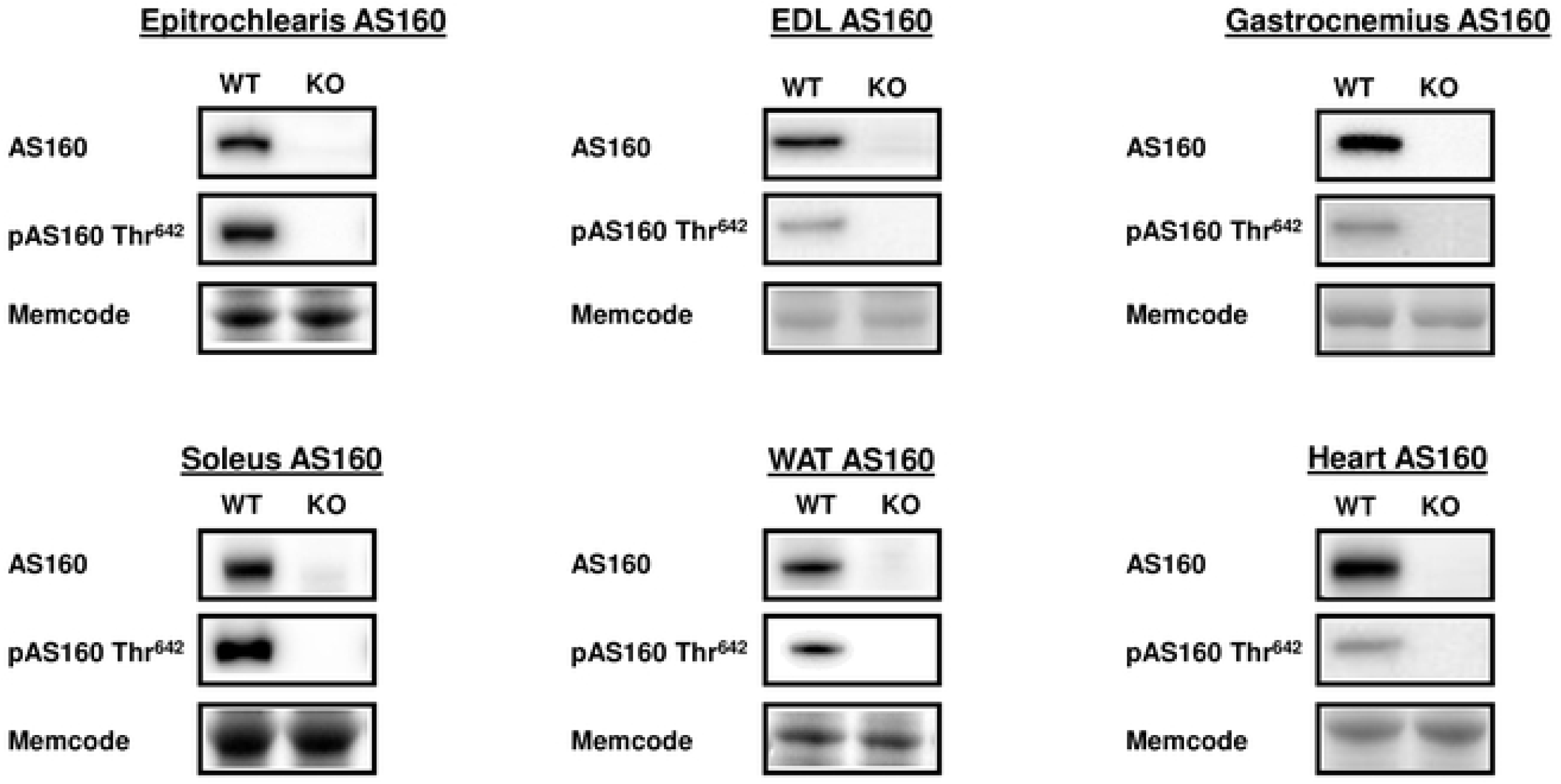
Tissue AS160 abundance and phosphorylation (pAS160 Thr^642^) from WT and KO rats subjected to the HEC. Extensor digitorum longus = EDL. White adipose tissue = WAT. Representative immunoblots and Memcode protein stain loading controls are included. AS160 abundance and pAS160 Thr^642^ were undetectable in all of the tissues from KO animals. N = 4 animals in each group.

### Body mass, body composition, and tissue masses

AS160-KO and WT rats were not different with regard to body mass, body composition (lean, fat, fluid) or tissue masses (Table 1). The ratios for skeletal muscle/body mass and heart mass/body mass did not differ for AS160-KO compared to WT rats.

**Table 1.** Body mass, body composition and tissue masses.

|  | **WT** | **KO** |
| --- | --- | --- |
| **Body Mass, g** | 183.75 ± 6.87 | 181.32 ± 5.88 |
| **Body Fat Mass, %** | 6.26 ± 0.26 | 6.66 ± 0.47 |
| **Body Lean Mass, %** | 75.29 ± 0.28 | 74.65 ± 0.52 |
| **Body Fluid Mass, %** | 7.18 ± 0.09 | 7.40 ± 0.17 |
| **Extensor digitorum longus, mg** | 107.98 ± 4.35 | 98.48 ± 6.72 |
| **Epitrochlearis, mg** | 45.42 ± 5.16 | 52.55 ± 4.60 |
| **Gastrocnemius, mg** | 1208.88 ± 36.80 | 1122.12 ± 46.37 |
| **Soleus, mg** | 127.55 ± 5.05 | 123.08 ± 5.68 |
| **Heart, mg** | 749.23 ± 22.77 | 750.08 ± 17.99 |

Body composition was measured for 7-8 weeks-old rats. The masses of skeletal muscles and hearts were measured for 11-weeks-old rats. Means ± SEM for 6 rats of each genotype: WT = wildtype, and KO = AS160-KO.

### Myosin heavy chain (MHC) isoform expression

Muscle fiber type composition based on relative MHC isoform abundance did not differ between the genotypes for epitrochlearis, soleus, EDL or gastrocnemius (Table 2).

**Table 2.**
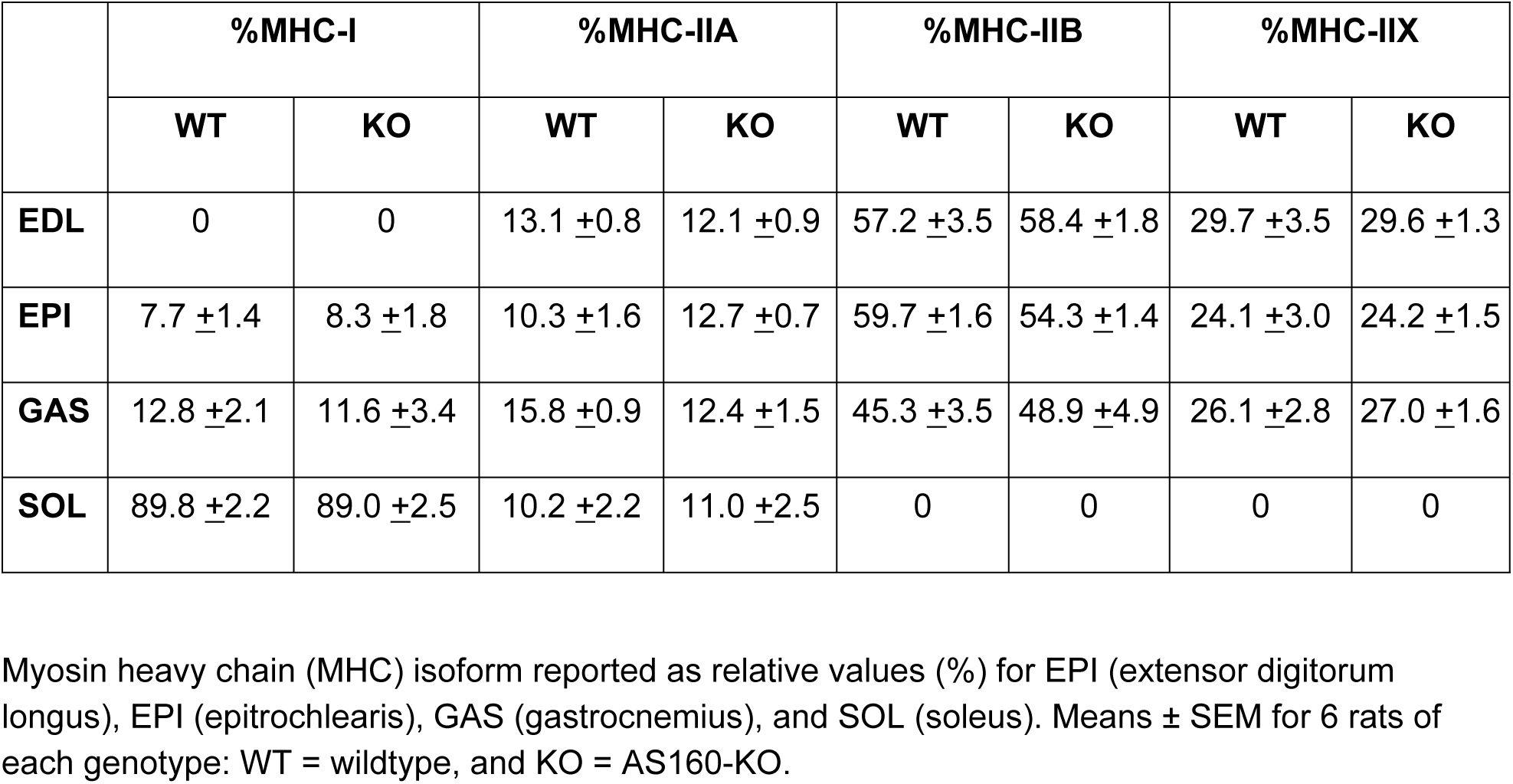
Relative myosin heavy chain isoform composition of skeletal muscles.

### Physical activity, indirect calorimetry, and food intake

There were no significant genotype-related differences for food intake (WT = 19.35 ±0.34 g/day; AS160-KO = 18.99 ±0.52 g/day), physical activity (total X-activity, ambulatory X-activity or vertical Z-physical activity), energy expenditure, oxygen consumption, respiratory exchange ratio (RER) (Table 3).

**Table 3.** Indirect calorimetry and physical activity.

|  | Dark |  | Light |  |
| --- | --- | --- | --- | --- |
|  | WT | AS160-KO | WT | AS160-KO |
| **VO<sub>2</sub> (ml/kg/h)** | 1987.9 ± 64.4 | 2113.3 ± 50.5 | 1554.8 ± 59.2 | 1643.3 ± 35.5 |
| **RER (VCO<sub>2</sub>/VO<sub>2</sub>)** | 0.933 ± 0.016 | 0.930 ± 0.023 | 0.896 ± 0.021 | 0.910 ± 0.025 |
| **Energy Expenditure (kcal/kg/h)** | 9.812 ± 0.296 | 10.429 ± 0.145 | 7.576 ± 0.241 | 8.069 ± 0.132 |
| **Activity-X Total (counts/h)** | 1459.7 ± 76.5 | 1596.3 ± 138.9 | 519.8 ± 54.5 | 532.2 ± 31.7 |
| **Activity-X Ambulatory (counts/h)** | 599.1 ± 57.5 | 662.1 ± 77.2 | 217.9 ± 29.8 | 227.2 ± 13.9 |
| **Activity-Z (counts/h)** | 470.5 ± 50.5 | 594.1 ± 103.1 | 186.8 ± 44.7 | 164.2 ± 23.1 |

Oxygen consumption rate = VO<sub>2</sub>. Respiratory exchange ratio = RER. Carbon dioxide production rate = VCO<sub>2</sub>. Means ± SEM for 4-6 rats of each genotype: WT = wildtype, and KO = AS160-KO.

### Oral glucose tolerance test (OGTT)

Glucose levels at baseline were slightly (12.6%), but significantly (P<0.05) greater for WT compared to AS160-KO females (Fig 2A). AS160-KO animals versus WT animals were glucose intolerant as evidenced by higher glucose concentration during minutes 15, 30, 60 of the OGTT (P<0.005 to 0.001) and on greater glucose AUC (area under the curve) (P<0.001; Fig 2). Insulin levels at baseline were not significantly different between AS160-KO and WT rats (Fig 2B).

**Figure 2.**
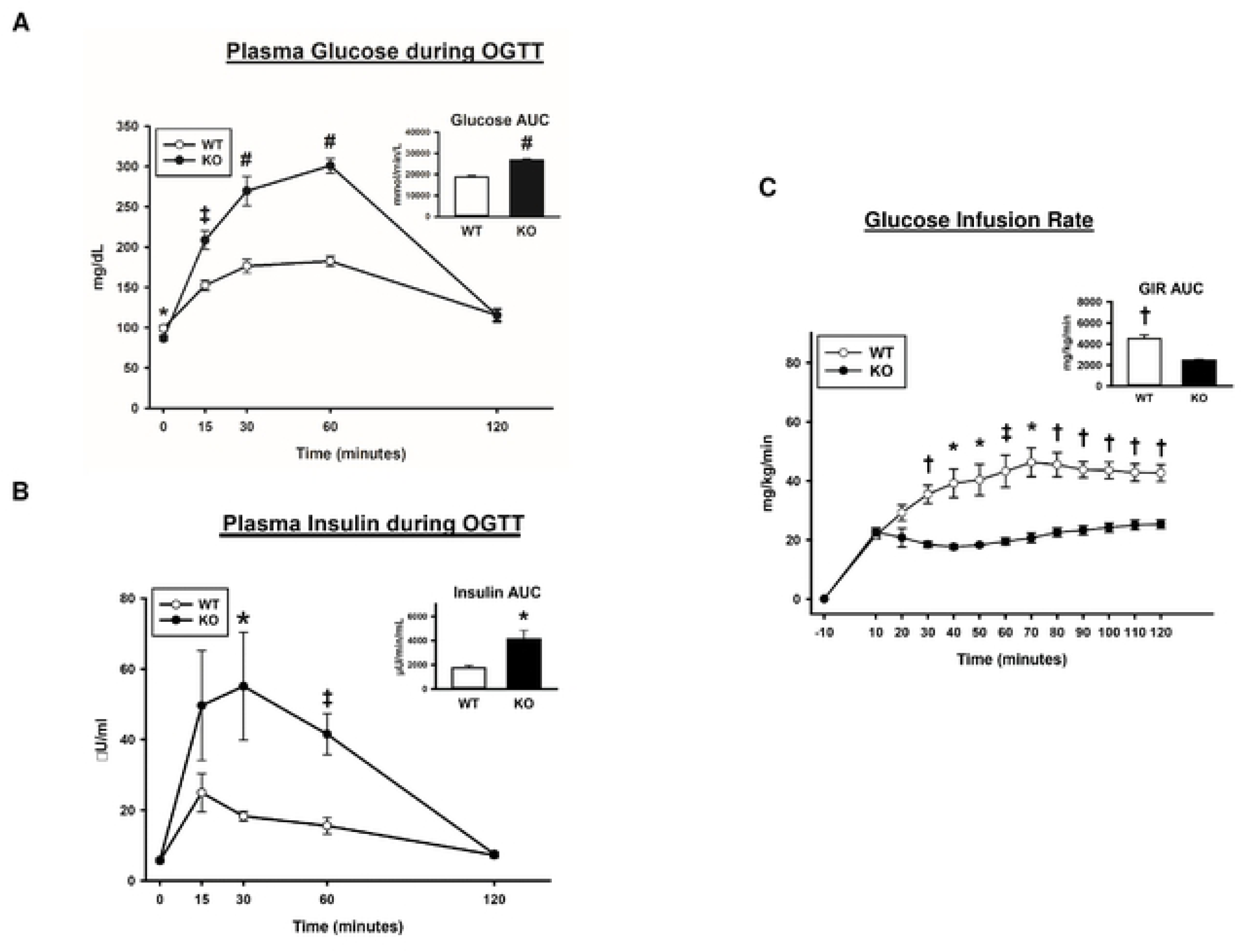
Oral glucose tolerance test = OGTT. Glucose infusion rate = GIR during the HEC in wildtype (WT; white bars and white circles) and AS160-KO (KO; black bars and black circles) rats. Area under the curve (AUC; inset) for (A) glucose and (B) insulin. Data were analyzed by Student’s t-test. *P<0.05, ^‡^P<0.005 and ^#^P<0.001 for WT versus KO rats. Means ± SEM for 6 rats of each genotype. (C) GIR for rats during the HEC. AUC for GIR (inset). Data were analyzed by Student’s t-test. *P<0.05, ^†^P<0.001 and ^‡^P<0.005 for WT versus KO. Means ± SEM for 4 rats of each genotype.

### Hyperinsulinemic-euglycemic clamp (HEC)

Although 6 WT and 6 AS160-KO rats underwent an HEC, 2 rats from each genotype were not successfully clamped because of technical problems. A catheter placed in one AS160-KO rat failed during the HEC, and the coefficient of variation (CV) for glycemia exceeded 20% for three rats (WT, n=2; AS160-KO n=1). Accordingly, the data from these 4 rats were not included in the statistical analysis of parameters dependent on blood glucose or insulin (Table 4), tissue glucose uptake and insulin signaling. The CV for the remaining 4 WT rats (3-14%) was comparable to the CV for the remaining 4 AS160-KO rats (5-12%), and the reported values for blood or plasma parameters, tissue glucose uptake, and phosphorylated signaling proteins from the HEC experiment represent the data collected from these 8 animals. No significant differences between AS160-KO compared to WT rats was found for blood glucose values, plasma insulin values, glucose turnover rate (GTR) or hepatic glucose production (HGP; Table 4). Blood glucose during the HEC was ∼18.6% lower (P<0.05) in AS160-KO compared to WT rats. Plasma insulin levels during the HEC did not differ for AS160-KO versus WT rats (Table 4). HGP and insulin suppression of HGP were not significantly different for AS160-KO compared to WT animals. The GIR (glucose infusion rate) AUC during the HEC was much lower for AS160-KO compared to WT rats (47.5%; P<0.001; Fig 2C). GIR was significantly lower at minutes 30, 40, 50, 60, 70, 80, 90, 100, 110, 120 (P<0.05 to 0.001). GTR was much less in AS160-KO compared to WT animals (65%; P<0.001). There were no differences between genotypes for plasma non-esterified fatty acid (NEFA) at baseline or during the HEC (WT versus AS160-KO: - 10 minutes, 0.489 ±0.014 versus 0.550 ±0.048; 80 minutes, 0.141 ±0.002 versus 0.152 ±0.005; 90 minutes, 0.131 ±0.006 versus 0.155 ±0.009; 120 minutes, 0.141 ±0.011 versus 0.16 ±0.011).

**Table 4.** Hyperinsulinemic euglycemic clamp (HEC)

|  | **WT** | **KO** |
| --- | --- | --- |
| **Basal blood glucose, mmol/L** | 6.81 ±0.41 | 6.13 ±0.24 |
| **Clamp blood glucose, mmol/L** | 7.49 ±0.43* | 6.10 ±0.15 |
| **Basal plasma insulin, µU/mL** | 7.21 ±1.65 | 6.75 ±0.54 |
| **Clamp plasma insulin, µU/mL** | 502.07 ±23.93 | 500.42 ±16.00 |
| **GIR, µmol/kg/min** | 240.02 ±13.27† | 135.87 ±6.98 |
| **Clamp GTR, µmol/kg/min** | 248.06 ±3.15† | 162.12 ±12.65 |
| **Basal HGP, µmol/kg/min** | 68.45 ±12.84 | 57.07 ±5.52 |
| **Clamp HGP, µmol/kg/min** | 8.03 ±10.88 | 26.24 ±7.32 |
| **Suppression of HGP rate, %** | 84.93 ±20.86 | 69.13 ±11.04 |

Glucose infusion rate = GIR. Glucose turnover rate = GTR. Hepatic glucose production = HGP. Means ± SEM for 4 rats of each genotype: WT = wildtype, and KO = AS160-KO. \*P 0.05, †P<0.001, WT versus KO.

### Glucose uptake by tissues during the HEC

In vivo glucose uptake was lower in the EDL (P<0.01) and the epitrochlearis (P<0.05) of AS160-KO versus WT rats (Fig 3). No significant genotype-related differences were observed in the soleus, gastrocnemius or white adipose tissue (WAT). AS160-KO versus WT rats had ∼3-fold greater glucose uptake by the heart (P<0.05).

**Figure 3.**
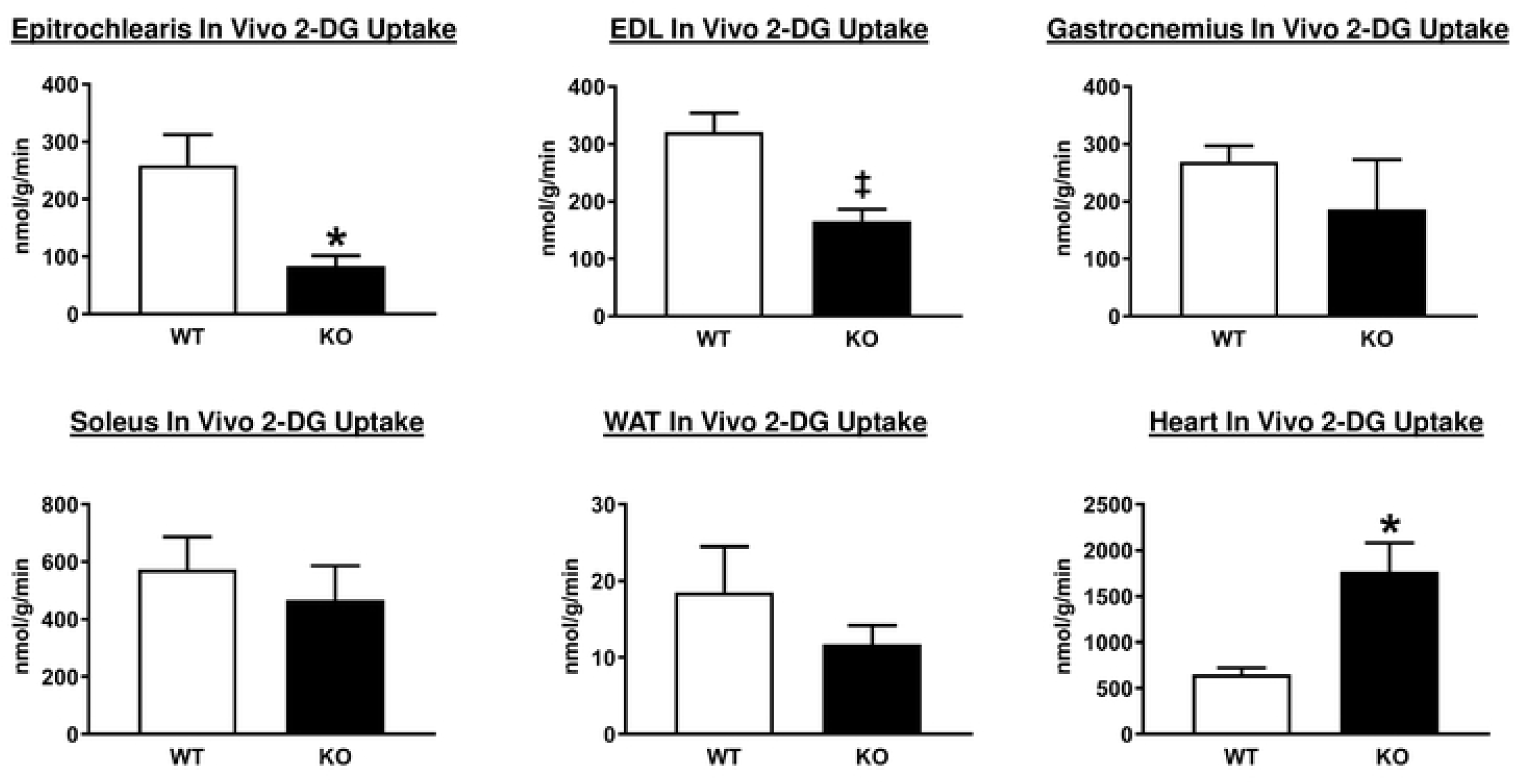
Tissue 2-deoxyglucose (2-DG) uptake in wildtype (WT; white bars) and AS160-KO (KO; black bars) rats subjected to the HEC. Extensor digitorum longus = EDL. White adipose tissue = WAT. Data were analyzed by Student’s t-test. *P<0.05, ^‡^P<0.01 for WT versus KO rats. Means ± SEM for 4 rats of each genotype.

### Immunoblotting of HEC tissues

AS160 and pAS160 Thr^642^ were both detected in tissues analyzed from WT rats and were not detectable in tissues analyzed from AS160-KO rats (Fig 1). Among all of the tissues evaluated, there were no significant genotype-related differences for either pAkt Ser^473^ or pAkt Thr^308^ (Fig 4). In addition, genotype-related differences were not found for TBC1D1, an AS160 paralog (Fig 5).

**Figure 4.**
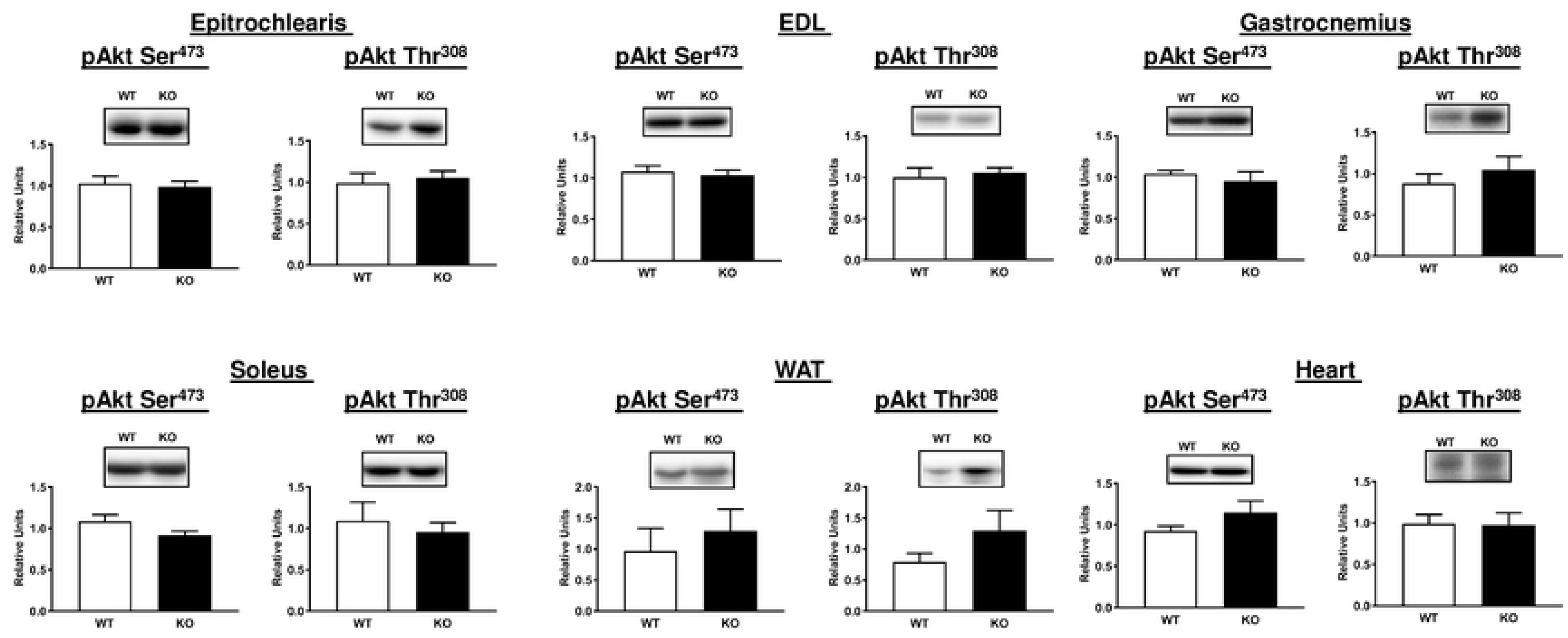
Akt phosphorylation on Ser^473^ (pAkt Ser^473^) and Thr^308^ (pAkt Thr^308^) in tissues from WT (white bars) and KO (black bars) animals subjected to the HEC. Extensor digitorum longus = EDL. White adipose tissue = WAT. To improve the clarity of these figures loading controls (Memcode protein stain) are not included on this figure or on subsequent figures (no genotype-related differences were found for any of the loading controls). Means ± SEM for 4 rats of each genotype.

**Figure 5.**
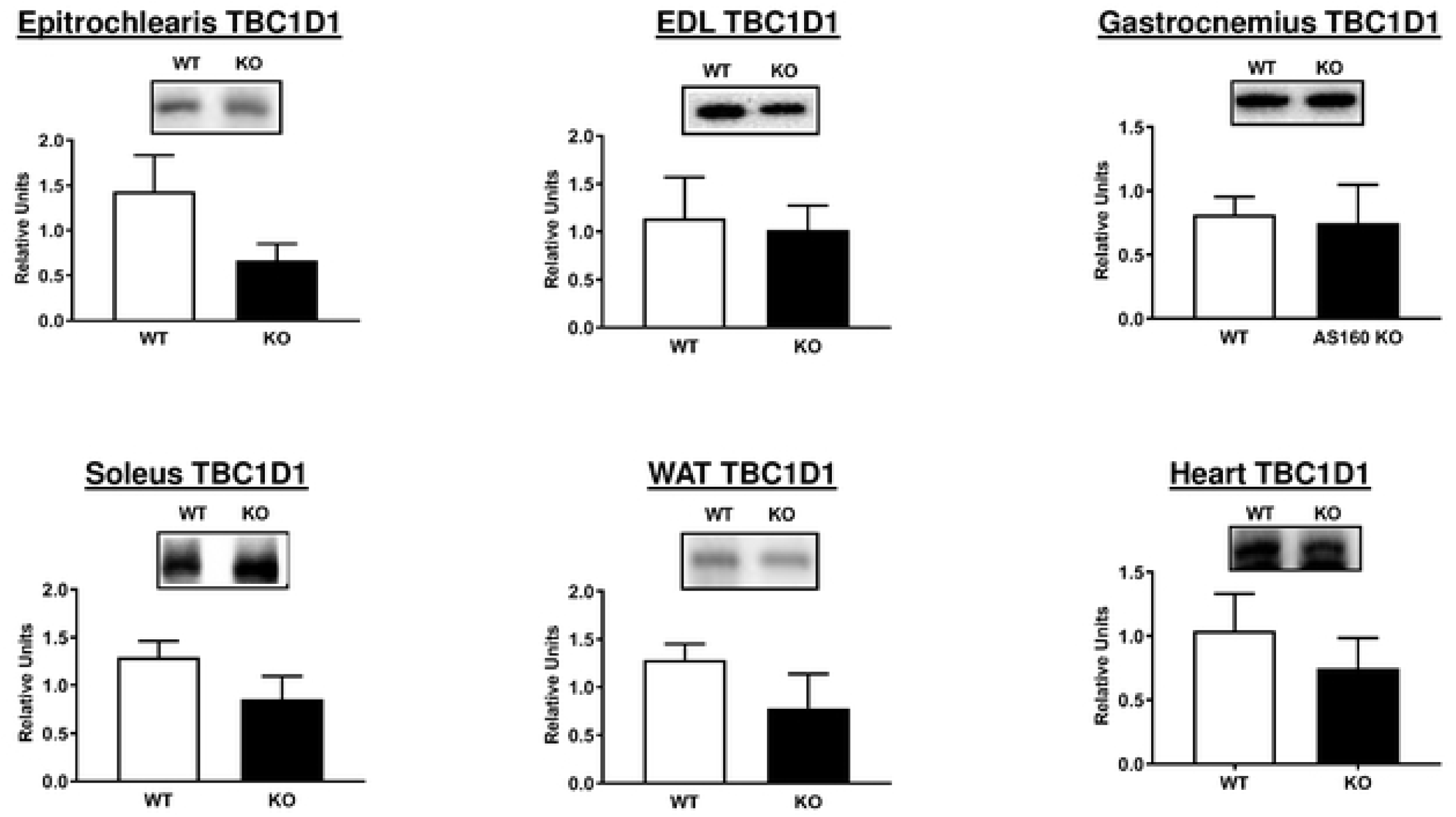
TBC1D1 abundance in tissues from WT (white bars) and KO (black bars) animals subjected to the HEC. Extensor digitorum longus = EDL. White adipose tissue = WAT. Means ± SEM for 4 rats of each genotype.

GLUT4 glucose transporter protein abundance was determined in each of the tissues after HEC. GLUT4 protein levels were much less in AS160-KO versus WT rats in the gastrocnemius, EDL, epitrochlearis, soleus, heart and WAT (Figure 6). GLUT1 glucose transporter protein content was not different between genotypes in any of these tissues (Supplemental Fig 1). In addition, hexokinase II abundance was not different between genotypes for any of these tissues (Supplemental Fig 2).

**Figure 6.**
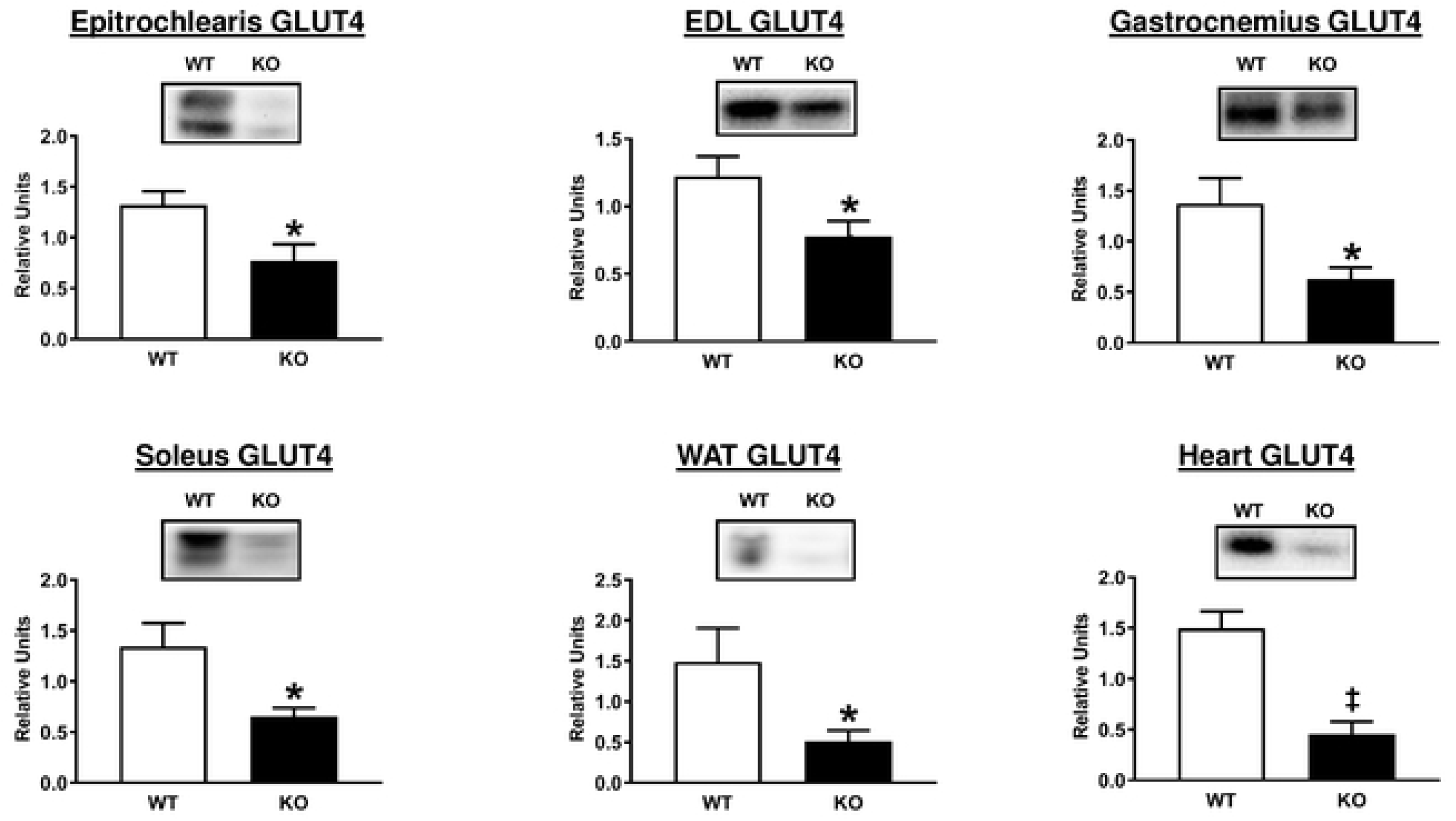
GLUT4 abundance in tissues from WT (white bars) and KO (black bars) animals subjected to the HEC. Extensor digitorum longus = EDL. White adipose tissue = WAT. *P<0.05, ^‡^P<0.01 for WT versus KO rats. Means ± SEM for 6 rats of each genotype.

Because Akt2 is required for insulin-stimulated glucose uptake by the heart [33], we determined Ser^474^ phosphorylation (pAkt2^Ser474^) in the heart from the HEC rats and found no difference between the AS160-KO and WT rats (Fig 7). Greater heart sodium-dependent glucose cotransporter 1 (SGLT1) expression has been found in diabetic animals concomitant with attenuated GLUT1 and GLUT4 levels [34], but we observed SGLT1 protein content did not differ between groups (Fig 7). Calcium ATPase (SERCA) has been linked to greater glucose uptake by the heart [35]. The abundance of myocardial SERCA2 was not different between genotypes (Fig 7). It was possible that the increased glucose uptake in AS160-KO heart was related to the reduced metabolism of another energy source, e.g., fatty acids. Therefore, we analyzed the abundance of the CD36 fatty acid translocase protein that is important for myocardial fatty acid uptake [36]. No significant genotype-related difference was observed for CD36 levels in the heart (Fig 7).

**Figure 7.**
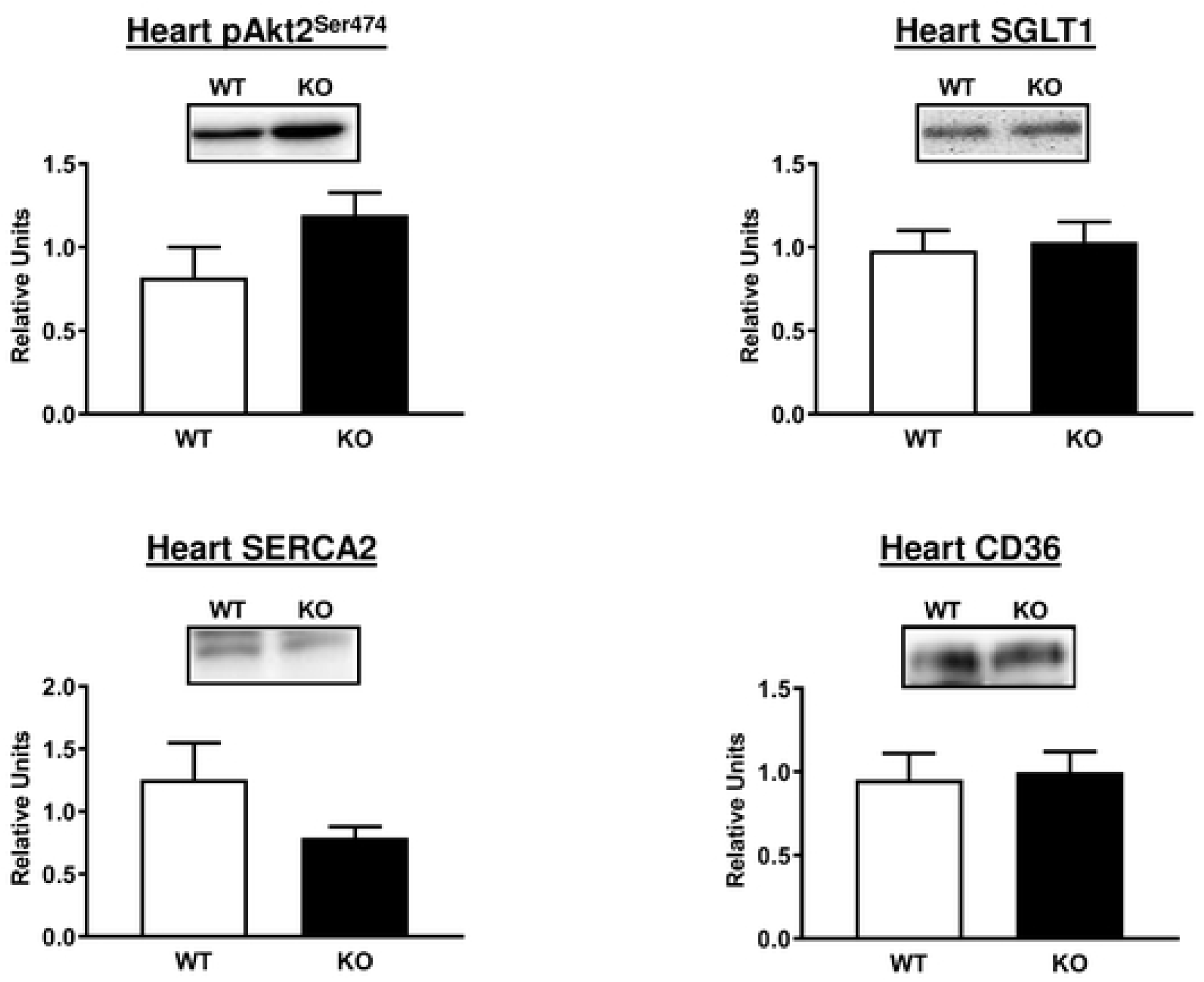
Akt2 phosphorylation on Ser^474^ (pAkt2^Ser474^) and abundance of SGLT1, SERCA2, and CD36 in heart from WT (white bars) and KO (black bars) animals subjected to the HEC. Means ± SEM for 4 rats of each genotype.

Activation of AMPK can lead to increased glucose uptake by the heart [37]. We determined the phosphorylation of AMPK at Thr172 (pAMPK Thr^172^), which is a major mechanism for increasing the enzyme’s activity [38]. We also assessed the phosphorylation of its substrate acetyl CoA-carboxylase (pACC Ser^79^) that is often used as an indicator for AMPK activity. However, no difference was detected between WT and AS160-KO for either pAMPK Thr^172^ or pACC Ser^79^ in the heart (Fig 8). Furthermore, phosphorylation of TBC1D1 on an AMPK-phosphosite, Ser237 (pTBC1D1 Ser^237^), was also undistinguishable in the heart of WT compared to AS160-KO rats (Fig 8).

**Figure 8.**
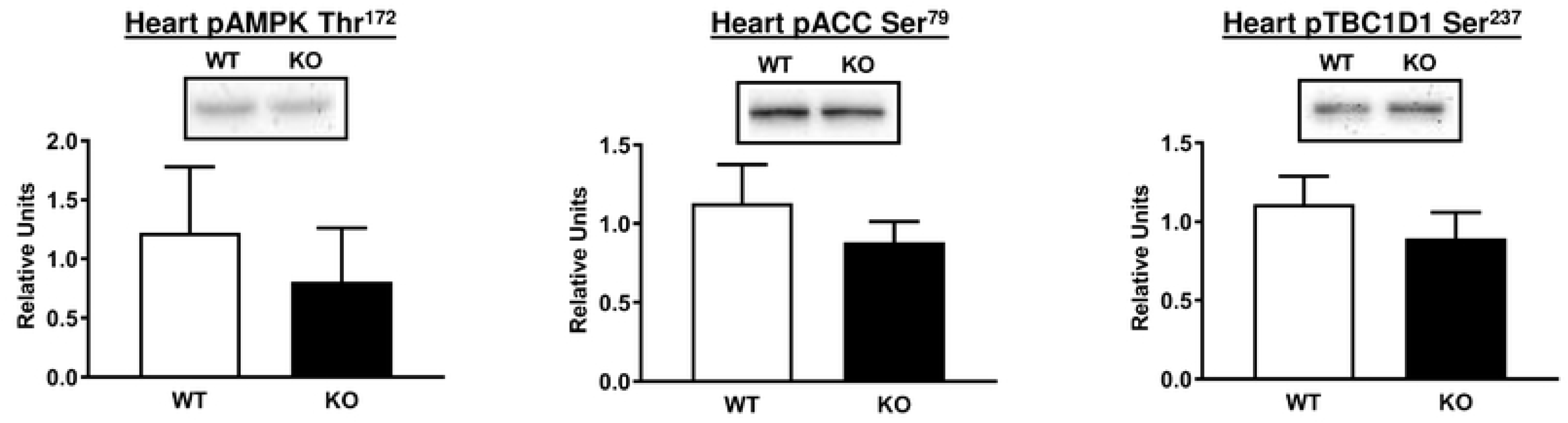
AMPK phosphorylation on Thr^172^ (pAMPK^Thr172^), phosphorylation of ACC on Ser^79^ (pACC Ser^79^) and phosphorylation of TBC1D1 Ser^237^ in heart from WT (white bars) and KO (black bars) animals subjected to the HEC. Means ± SEM for 4 rats of each genotype.

We also assessed the abundance of GLUT8 glucose transporter protein, which is known to be expressed by the heart [39]. However, no genotype-related difference was found for myocardial GLUT8 abundance (Fig 9). Because major metabolic fates of glucose include conversion to lactate or mitochondrial oxidation, we also determined the abundance of LDH and multiple components of the electron transport chain and oxidative phosphorylation (NDUFB8, SDHB, UQCRC2, MTCO1, and ATP5A). No differences between WT and AS160-KO rats were detected for LDH (Fig 9) or any of the mitochondrial proteins that were studied (Supplemental Fig 3).

**Figure 9.**
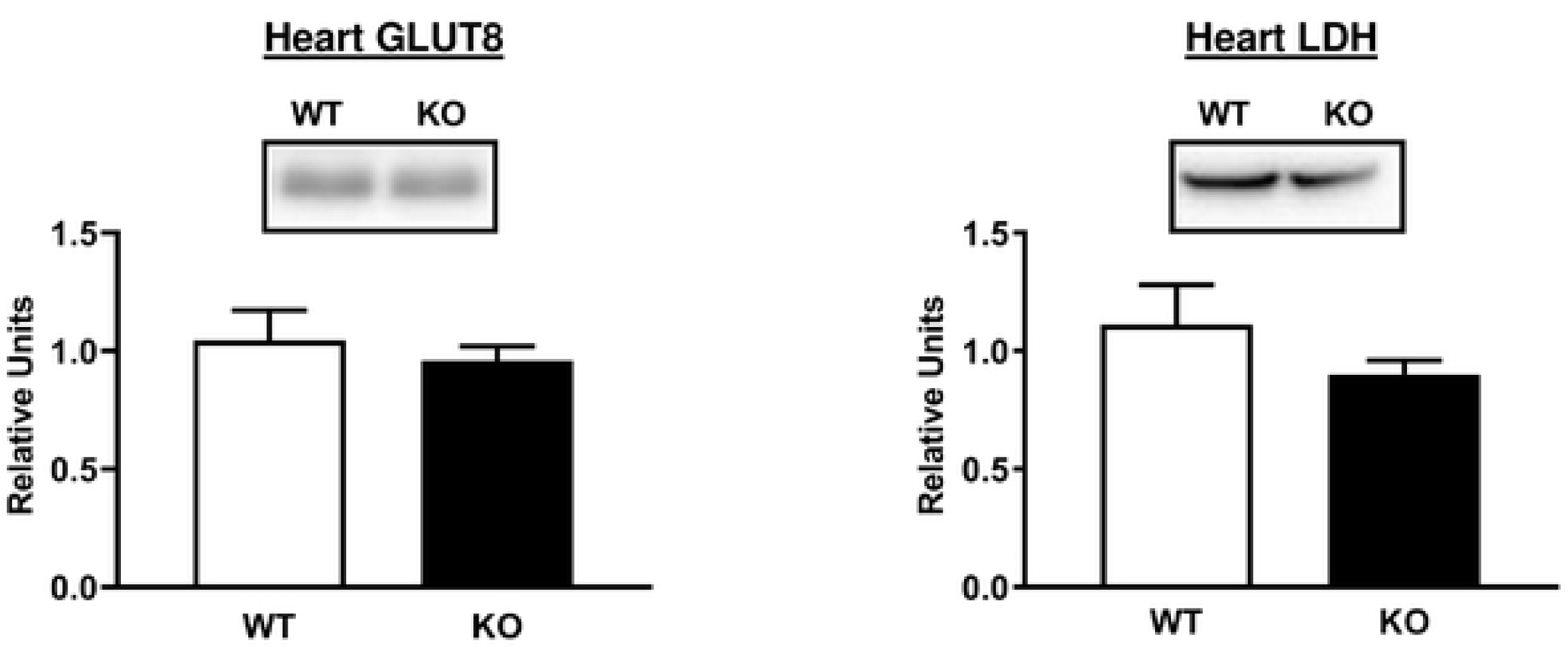
GLUT8 and LDH abundance in tissues from WT (white bars) and KO (black bars) animals subjected to the HEC. Means ± SEM for 4 rats of each genotype.

### Glucose uptake and immunoblotting of ex vivo insulin-stimulated skeletal muscle

Soleus and epitrochlearis muscles were studied ex vivo with and without an insulin dose (500 µU/ml) that corresponded to the plasma insulin values that the animals were exposed to during the HEC. Glucose uptake by insulin-stimulated soleus (P<0.001) and epitrochlearis (P<0.001) muscles isolated from WT rats was much greater than values in AS160-KO rats (Fig 8). Glucose uptake by the epitrochlearis from AS160-KO animals was greater for insulin-stimulated compared to paired muscles without insulin (P<0.001). Glucose uptake with insulin was increased compared to without insulin in the soleus (P<0.05) and the epitrochlearis (P=0.05) from AS160-KO rats. Glucose uptake in the epitrochlearis without insulin was significantly greater (P<0.05) for WT versus AS160-KO rats (Fig 10).

**Figure 10.**
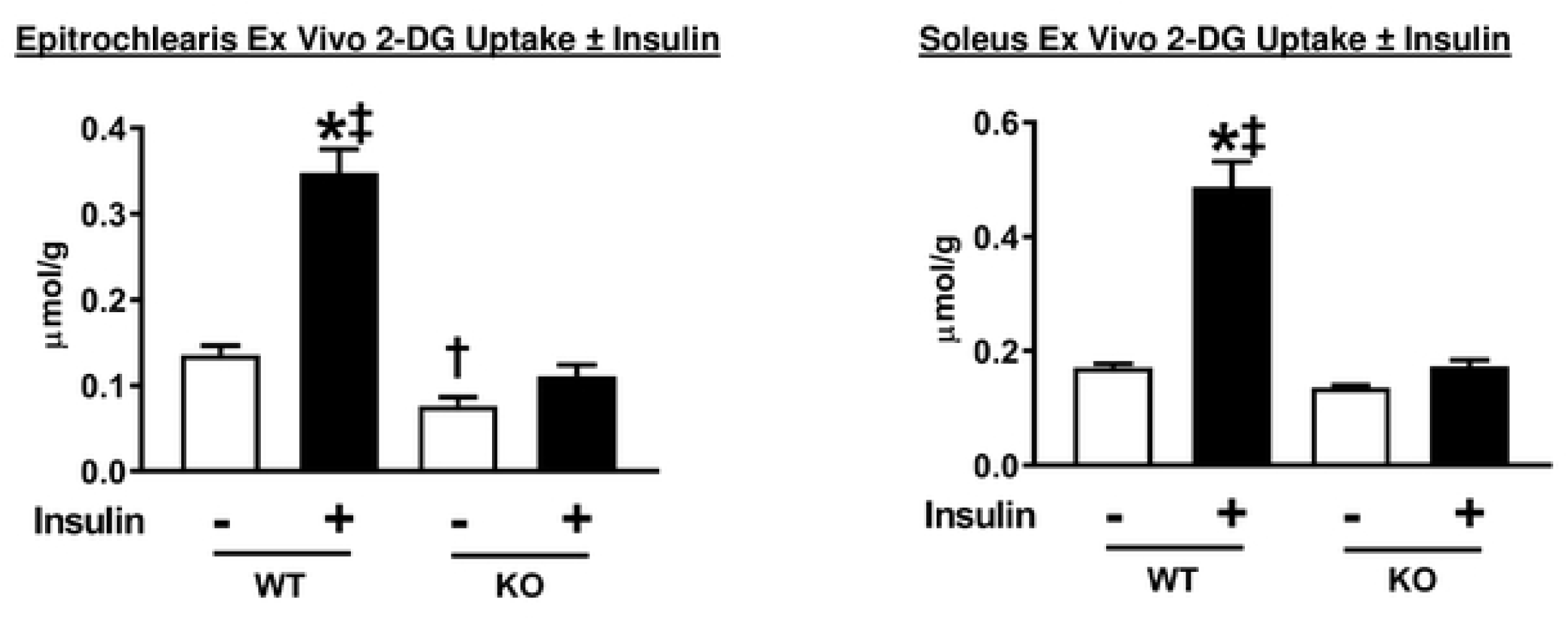
2-Deoxyglucose (2-DG) uptake by isolated epitrochlearis and soleus muscles from WT and KO rats. Paired muscles were incubated in the absence (white bars) or presence (black bars) of insulin (500 µU/ml). Data were analyzed by two-way ANOVA, and Holm-Sidak post hoc analysis was used to identify the source of significant variance. *P<0.001, without insulin versus insulin of the same genotype; ^‡^P<0.001, WT versus KO with the same insulin dose; ^†^P<0.05, WT versus KO with the same insulin dose. A t-test revealed significantly greater glucose uptake with insulin versus without muscle in soleus (P<0.05). A non-significant trend (P=0.05) for an insulin-stimulated increase was found with a t-test in epitrochlearis. Means ± SEM for 8-9 rats of each genotype.

pAS160 Thr^642^ in soleus (P<0.01) and epitrochlearis (P<0.01) muscles from WT rats was greater with insulin versus without insulin (Fig 11). pAS160 Thr^642^ was undetectable in either epitrochlearis or soleus muscles from AS160-KO rats (Figure 11 A-B). With insulin treatment, a substantial increase was found for the phosphorylation of Akt Ser^473^ and Thr^308^ (Fig 11 C-F) in the soleus and epitrochlearis. Insulin-stimulated phosphorylation of Akt was not different between genotypes in either the soleus or epitrochlaris.

**Figure 11.**
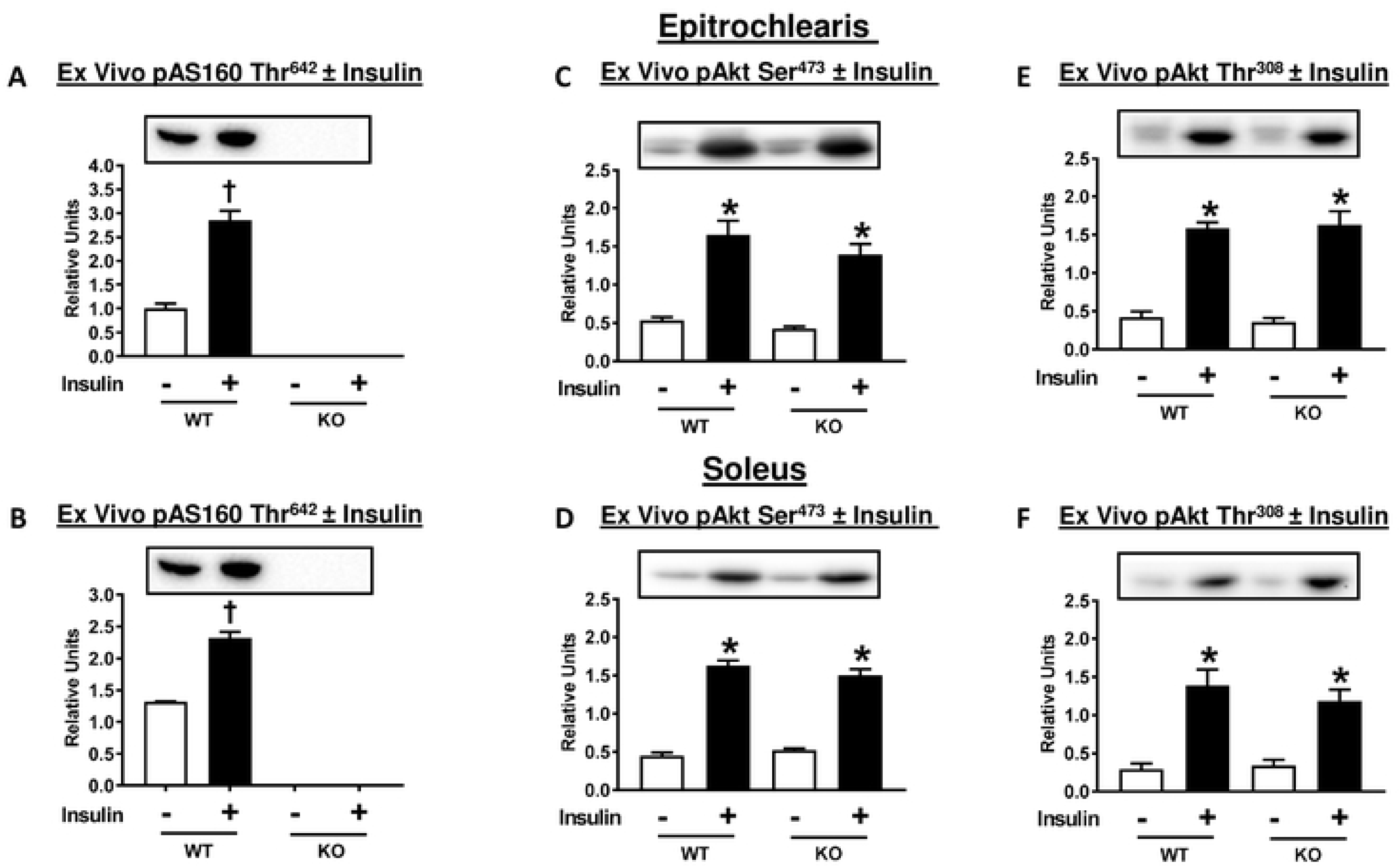
Phosphorylation of AS160 (pAS160 Thr^642^) and Akt (pAkt Ser^473^ and pAkt Thr^308^) for isolated epitrochlearis and soleus muscles from WT and KO rats. Paired muscles were incubated in the absence (white bars) or presence (black bars) of insulin (500 µU/ml). (A) and (B), AS160 pThr^642^ was not detectable in any of the tissues studied from KO rats. Data were analyzed by t-test. ^†^P<0.01, no insulin versus insulin in WT rats. Means ± SEM for 3 rats of each genotype. (C), (D), (E), and (F), * P<0.001, no insulin versus insulin in rats with same genotype. Means ± SEM for 6 rats of each genotype.

### Glucose uptake and immunoblotting of ex vivo AICAR-stimulated skeletal muscle

AICAR effects on soleus glucose uptake was not determined because earlier results indicated that AICAR does not alter glucose uptake by rat soleus [28]. Glucose uptake by the epitrochlearis from WT rats was significantly (P<0.001) greater with AICAR versus without AICAR (Fig 12A). WT versus AS160-KO rats had much higher AICAR-stimulated glucose uptake (P<0.001). In AS160-KO rats, a paired *t*-test indicated that glucose uptake by muscles incubated in the presence of AICAR compared to muscles incubated in the absence of AICAR were not significantly different (P<0.15).

**Figure 12.**
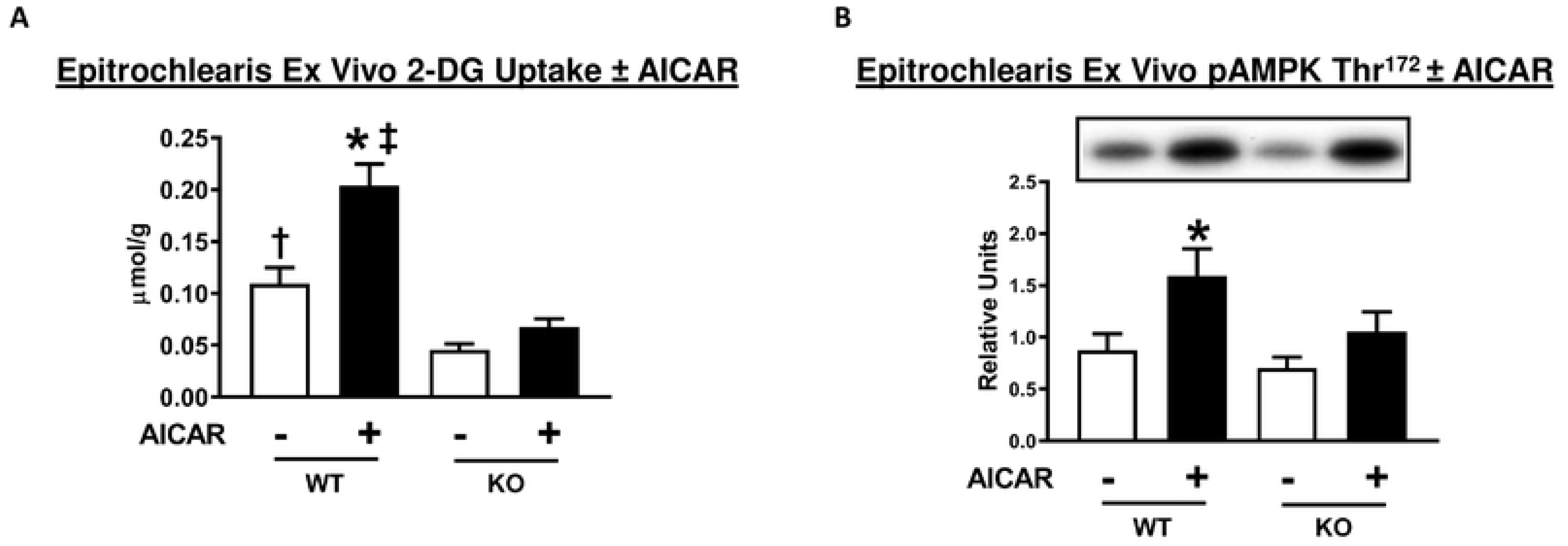
2-Deoxyglucose (2-DG) uptake and AMP-activated protein kinase (AMPK) phosphorylation (pAMPK Thr^172^) in isolated epitrochlearis from WT and KO rats. Paired muscles were incubated in the absence of (white bars) or presence of (black bars) AICAR (2 mM). Data were analyzed by two-way ANOVA, and the Holm-Sidak post hoc test. (A) *P<0.001 for no AICAR versus AICAR in the same genotype; ^‡^P<0.001, WT versus KO with the same AICAR dose; ^†^P<0.05 for WT versus KO without AICAR. Means ± SEM for 5-9 rats of each genotype. (B) *P<0.05 for no AICAR versus AICAR within the same genotype. A t-test revealed that pAMPK Thr^172^ was significantly greater (P<0.05) for muscles incubated with AICAR versus without AICAR in KO rats. Means ± SEM for 6-9 rats of each genotype.

As expected, in the muscles from WT rats, AICAR treatment resulted in significantly (P<0.05) greater pAMPK Thr^172^ compared to muscles without AICAR treatment (Fig 12B). There was no difference between WT and AS160-KO groups for pAMPK Thr^172^. AICAR-treatment significantly (P<0.05) increased pAMPK Thr^172^ in muscles from AS160-KO rats based on a paired student t-test.

## Discussion

The only previously published research on AS160-KO rats focused exclusively on male rats [5]. Therefore, the current study filled an important gap in knowledge by comparing a large number of metabolic outcomes in female AS160-KO versus WT rats. There has also been a marked disparity in the amount of data previously reported using female compared to male AS160-KO mice. Much of the published research on AS160-KO mice has either entirely or largely focused on males [4, 7, 8], with only two earlier studies on AS160-KO mice including data for some, but not all, endpoints in both sexes [3, 9]. Published research on AS160-deficient humans has not separately reported data for males and females [6].

The current results revealed that AS160 deficiency had metabolic consequences for female rats that were comparable to the recently reported results for male AS160-KO rats [5]. Similar to males, the female AS160-KO versus WT rats were glucose intolerant based on the OGTT and insulin resistant based on GIR and GTR during the HEC. Insulin resistance in female AS160-KO rats during the HEC was largely attributable to lower glucose uptake by skeletal muscles, including the epitrochlearis and EDL. GLUT4 glucose transporter protein abundance was substantially lower for female AS160-KO compared to WT rats in each of the skeletal muscles studied (epitrochlearis, EDL, soleus, and gastrocnemius). In male rats, a significant genotype-related deficit in GLUT4 was previously reported for the epitrochlearis, EDL and soleus, and there was a non-significant trend for lower GLUT4 content in the gastrocnemius [5]. Among the dozens of measurements made in both male and female rats, gastrocnemius GLUT4 was the only endpoint that had a significant difference in one sex, but not in the other. The insulin resistance in skeletal muscle was not attributable to genotype differences for GLUT1 or HKII abundance, Akt phosphorylation or fiber type (based on myosin heavy chain isoform expression) in female rats. Similarly, none of these endpoints differed between male AS160-KO and WT rats [5].

Analysis of glucose uptake by isolated skeletal muscles is useful because it provides information about the tissue’s intrinsic ability for glucose uptake. Both the epitrochlearis and soleus muscles from female AS160-KO rats were profoundly insulin resistant, similar to the results for male AS160-KO rats [5]. Earlier results from male AS160-KO rats [5] and mice [7] demonstrated lower glucose uptake compared to WT controls in response to stimulation with the AMPK-activator AICAR. The reduction in AICAR-stimulated glucose uptake by the epitrochlearis of either female or male AS160-KO versus WT rats indicates that the insulin resistance was not specific to insulin-stimulation. This result supports the idea that reduced glucose uptake with either insulin or AICAR is likely secondary, at least in part, to lower GLUT4 protein levels.

The greatest genotype-difference in the female rats was the markedly greater myocardial glucose uptake rate in the AS160-KO compared to WT rats. We previously reported an almost identical relative increase in heart glucose uptake in male AS160-KO versus WT rats [5]. In both sexes, the difference in myocardial glucose uptake occurred in spite of much lower GLUT4 protein abundance in the AS160-KO compared to WT rats. In neither sex was the greater glucose uptake attributable to altered myocardial abundance of GLUT1, HKII or SGLT1. Furthermore, in the females, we also assessed heart GLUT8, and found that the abundance of this glucose transporter was also unaltered by AS160 deficiency. Taking together all of these results, there is no evidence that the genotype effect on myocardial glucose uptake is linked to greater expression of glucose transporter proteins. Our working hypothesis is that the result is attributable to altered subcellular localization of glucose transporter proteins, with greater cell surface glucose transporter content in the AS160-KO compared to WT rats of either sex. GLUT4 is the most abundant glucose transporter protein expressed by the heart, and GLUT4 translocation is normally the dominant mechanism for increasing myocardial glucose uptake [40]. Therefore, even though AS160 deficiency was characterized by a substantial decline in total GLUT4 abundance in the heart, increased GLUT4 translocation may be hypothesized to play a role in the observed 3-fold increase in myocardial glucose uptake.

AMP-activation has been reported to increase myocardial GLUT4 translocation and glucose uptake [41]. Accordingly, we evaluated the activation of AMPK phosphorylation on Thr172 because increased AMPK phosphorylation on this site results in a substantial increase in AMPK activity [38]. We also assessed the phosphorylation of AMPK’s substrate acetyl CoA-carboxylase (ACC) which is often used as a surrogate indicator of AMPK activation [42, 43]. In addition, we determined phosphorylation of the Rab-GTPase activating protein TBC1D1 on Ser231, an AMPK-phosphomotif that has been implicated in TBC1D1’s regulation of glucose uptake [44]. However, there were no differences between genotypes for phosphorylation of AMPK, ACC or TBC1D1, indicating that other mechanisms are responsible for the greater myocardial glucose uptake in AS160-KO versus WT rats.

We probed several other potential factors that might be linked to altered myocardial glucose metabolism. Cardiac hypertrophy and heart failure are characterized by greater dependence on glucose for energy [45, 46]. However, neither heart mass nor heart mass/body mass ratio differed between genotypes. The heart is normally able to switch between carbohydrate and lipid as a fuel source, and it seemed possible that differences in lipid metabolism might be relevant to the marked genotype-effect on glucose uptake by the heart. However, neither plasma NEFA concentration nor myocardial abundance of the fatty acid translocase CD36 were different for female AS160-KO versus WT rats. Myocardial SERCA overexpression has been reported to increase glucose uptake by the heart [35], but there was no evidence for greater SERCA2 abundance in the female AS160-KO rats. It seemed possible that the marked increase in glucose uptake by the heart might be related to modifications in the expression of key mitochondrial proteins or LDH. However, no genotype differences were detected for either of these parameters.

In contrast to the striking genotype-related differences in multiple glucoregulatory endpoints, the AS160-KO and WT rats were quite similar for many other characteristics. Body composition, energy expenditure and fuel selection, food intake, spontaneous physical activity, tissue masses, and skeletal muscle myosin heavy chain isoform composition did not differ between genotypes in the female rats. Similarly, no genotype-related differences were detected for any of these endpoints in male AS160-KO compared to WT rats [5]. Although many of these outcomes have been linked to altered glucoregulation and insulin sensitivity, none of the parameters were responsible for the marked genotype differences that were found in AS160-KO compared to WT rats of either sex.

In conclusion, the current results using female AS160-KO rats taken together with the findings of our recent study using male AS160-KO rats [5] revealed that the metabolic phenotypes of both sexes were comparably responsive to AS160 deficiency. The current study also further advanced earlier research by including novel analyses aimed at elucidating the mechanisms responsible for the substantial increase in myocardial glucose uptake of AS160-KO versus WT rats. The results provided evidence that this outcome was not attributable to greater abundance of multiple glucose transporter proteins or enhanced activation of the AMPK pathway. The insights from the current study provide useful knowledge for future research aimed at elucidating the roles that AS160 plays in controlling glucose uptake and other biological processes in female rats.

## Acknowledgments

The authors thank the staff of the University of Michigan Animal Phenotyping Core for services in the OGTT, HEC, indirect calorimetry, physical activity, plasma NEFA, and body composition analyses. We appreciate Dr. Swati Agrawal’s role in the initial generation of the CRISPR/Cas9 animals. We acknowledge Elizabeth Hughes for CRISPR/Cas9 target design and construction of genome editing reagents, Wanda Filipiak for rat zygote microinjection and the Transgenic Animal Model Core of the University of Michigan’s Biomedical Research Core Facilities. The authors thank Dr. Jeremie Ferey from Washington University School of Medicine in Saint Louis for generously providing the GLUT8 antibody.

## Author contributions

**Conceptualization:** Gregory D. Cartee

**Data curation:** Xiaohua Zheng, Edward B. Arias, Gregory D. Cartee

**Formal analysis:** Xiaohua Zheng, Edward B. Arias, Nathan R. Qi

**Funding acquisition:** Gregory D. Cartee

**Investigation:** Edward B. Arias, Xiaohua Zheng, Nathan R. Qi

**Methodology:** Gregory D. Cartee, Thomas L. Saunders

**Supervision:** Gregory D. Cartee

**Validation:** Xiaohua Zheng, Edward B. Arias

**Visualization:** Xiaohua Zheng, Edward B. Arias

**Writing – original draft:** Xiaohua Zheng, Edward B. Arias, Gregory D. Cartee

**Writing – review & editing:** Xiaohua Zheng, Edward B. Arias, Nathan R. Qi, Thomas L. Saunders, Gregory D. Cartee

